# Distance Measures for Tumor Evolutionary Trees

**DOI:** 10.1101/591107

**Authors:** Zach DiNardo, Kiran Tomlinson, Anna Ritz, Layla Oesper

**Affiliations:** Department of Computer Science, Carleton College, Northfield, MN, USA.; Department of Biology, Reed College, Portland, OR, USA.

## Abstract

In recent years, there has been increased interest in studying cancer by using algorithmic methods to infer the evolutionary tree underlying a tumor’s developmental history. Quantitative measures that compare such trees are then vital to benchmarking these algorithmic tree inference methods, understanding the structure of the space of possible trees for a given dataset, and clustering together similar trees in order to evaluate inheritance patterns. However, few appropriate distance measures exist, and those that do exist have low resolution for differentiating trees or do not fully account for the complex relationship between tree topology and how the mutations that label that topology are inherited. Here we present two novel distance measures, **C**ommon **A**ncestor **Set** distance (CASet) and **D**istinctly **I**nherited **S**et **C**omparison distance (DISC), that are specifically designed to account for the subclonal mutation inheritance patterns characteristic of tumor evolutionary trees. We apply CASet and DISC to two simulated and two breast cancer datasets and show that our distance measures allow for more nuanced and accurate delineation between tumor evolutionary trees than existing distance measures. Implementations of CASet and DISC are available at: https://bitbucket.org/oesperlab/stereodist.

## 1 Introduction

A tumor is the result of an evolutionary process and its history can therefore be described as a rooted tree [1]. Specifically, the tree’s root can either represent a healthy cell or the original founding tumor population (assuming monoclonal tumor evolution), and every other vertex represents a distinct tumor population that existed at some point during the tumor’s evolution. Directed edges represent direct ancestral relationships between populations. In recent years, a number of methods have been developed to infer the tree describing a tumor’s evolution from single nucleotide variants in bulk sequencing data [2–9] and single cell sequencing data [10–14] (see [15] for a more complete listing). The goal of tree inference is to gain a better understanding of tumor development, which may in turn reveal insights about the mutations that drive a tumor’s growth [16, 17] and may be targeted for patient treatment [18, 19].

Due to the ongoing development of tumor evolution inference methods, the similarity of two potential tumor histories often needs to be quantified. First, new methods need to be benchmarked against other methods or against a ground truth tree, and the ad hoc measures that have typically been used in these situations [2–4] have not been rigorously studied. Second, some methods themselves rely on the use of a distance measure when inferring a tumor’s evolutionary history. For example, the GraPhyC method [20] uses a distance measure to create a consensus tumor history from several input histories. Lastly, there have been growing questions about the structure of the space of possible evolutionary histories consistent with the underlying sequence data [21–23] and how tumor evolutionary histories across patients can be used to identify patterns of tumor evolution [24, 25]. Further analysis of these questions would be aided by distance measures tuned to the intricacies of tumor evolution histories.

For traditional phylogenetic trees, there are several well-known distance measures such as Robinson-Foulds [26] and triplet distance [27], among others. However, the substantial evolutionary differences between species and cancer subpopulations prevent the direct application of traditional phylogenetic distance methods [15]. The primary difference between the two types of trees is that tumor evolutionary trees have internal node labels, which represent mutations rather than extant species. Furthermore, the mutations in a tumor evolutionary tree are inherited by all descendant tumor populations, creating a complex underlying relationship between topology and mutation labeling. In addition, nodes in tumor evolutionary trees may contain multiple labels (indicating mutations whose order of appearance cannot be readily identified) whereas nodes in a traditional phylogeny contain only one label each [20, 28]. There is then a critical need for specially calibrated distance methods that account for the intricacies of tumor evolutionary trees.

Despite this need for tumor tree distance measures, a limited number of such measures have been rigorously developed and comprehensively evaluated. Several simple distance measures described by [20] generalized earlier ad hoc approaches that relied on the existence of a ground truth tree [2–4], but were not the focus of that work, and their effectiveness was not analyzed in depth. Another recently proposed approach called MLTED uses an edit distance-based measure focused on handling multi-labeled nodes to count the minimum number of moves to convert both trees into a specific common tree [28]. However, this distance measure does not explicitly consider that mutation labels are inherited by all descendant populations. We argue that distance measures that consider how mutation placement affects a tree’s global structure will allow for much more precision when comparing tumor evolutionary trees.

In this work, we formalize the definition of a *tumor evolution distance measure* by precisely defining the input and output of such a function and describing what features of such a measure are desirable in the context of tumor evolution. We then describe two novel tumor evolution distance measures, CASet and DISC, that are specifically designed to account for structure of mutation inheritance by subsequent tumor populations and extend these measures to trees that do not share the same set of mutations. We apply our distance measures to two simulated datasets and two breast cancer datasets. We find that CASet and DISC allow for more precision when comparing tumor evolution histories and are better able to distinguish groups of similar trees than existing distance measures when applied in a clustering scenario.

## 2 Methods

### 2.1 Tumor Evolutionary Trees

We first make two common assumptions about tumor evolution. The first is the *infinite sites assumption* (ISA) which states that no mutation occurs more than once during a tumor’s history, and that once gained, a mutation is never lost. This has been a common assumption made by many methods that infer tumor evolutionary histories ([2–4, 6, 8] and many others). While some recent phylogeny inference methods do allow for minor violations of the ISA (e.g. [29]), our work here will be most widely applicable if we assume the ISA. The second assumption is that all tumor cells are descended from a single founding tumor cell, and hence the tumor’s evolution can be described as *monoclonal*. This assumption is non-essential to our approaches and can easily be dropped by rooting evolution trees with healthy cells instead of founding tumor mutations. Nonetheless, we make the monoclonal assumption to simplify our definitions. We now formally describe the evolutionary history of a monoclonal tumor adhering to the ISA as a clonal tree.

#### Definition 2.1.

*A **clonal tree** is a rooted, directed tree T in which: (i) each vertex in the tree is labeled by one or more mutations; and (ii) no mutation appears more than once*.

We also define ℑ to be the set of all clonal trees. Given a tree *T* ∈ ℑ, we define *M*(*T*) to be the set of all mutations (i.e. vertex labels) in *T*. In this representation, every vertex represents a distinct tumor clone (or population) that existed at some point during the tumor’s evolution. Directed edges represent direct ancestral relationships between tumor clones. The mutation labels indicate the clone in which the mutation first appeared. Thus, the complete set of mutations that exist in any particular clone, represented by vertex *v*, is the set of mutations that label all vertices on the path from the root to vertex *v*.

We sometimes wish to restrict our attention to a predetermined set of mutations, so we also define *m*-clonal trees for this purpose.

#### Definition 2.2.

*An **m-clonal tree** is a clonal tree T with M*(*T*) = {1, …, *m*}.

We emphasize that this definition uses the variable *m* to refer to the set of mutations rather than the number of clones in the tree. We also define ℑ_*m*_ to be the set of all *m*-clonal trees that share the same mutation set [*m*] = {1, 2, …, *m*}.

### 2.2 Tumor Evolution Distances

In this section, we define a tumor evolution distance measure on clonal trees and then analyze what particular features are desirable for such a measure.

#### Definition 2.3.

*A **tumor evolution distance measure** is a function d*: ℑ × ℑ → ℝ^*≥*0^ *for which a value of d*(*·, ·*) *that is close to 0 indicates that the two input clonal trees are very similar and progressively larger values of d*(*·, ·*) *indicate the clonal trees are more dissimilar.*

A tumor evolution distance measure must give us a quantitative evaluation of how different two tumor histories are from each other, but how best to define “different” is not immediately obvious. There are two main aspects of tumor evolutionary trees that should contribute to a distance measure: (i) the topology of the tree; and (ii) the labels present in the vertices of the trees. Topology can be separated from labeling by simply ignoring all labels, and the labels can be separated from topology by considering only the the set(s) of labels that appear or appear together. Thus, simple distance measures could certainly consider each of these aspects separately. However, these two tree attributes are inherently intertwined.

Since mutation labelings indicate in which vertex a mutation was first acquired, all descendants of that vertex also inherit that mutation. A difference in a vertex with many descendants should then contribute more to a distance measure than one in a vertex with few descendants, since it affects more clonal populations. Thus, a tumor evolution distance measure that simply counts the differences between trees (often referred to as a tree edit distance, as proposed by [28, 30], and others) does not address the impact any given label change may have. A distance measure should assign different weights to disagreements in different locations in order to appropriately address the relationship between topology and mutation labeling. These observations and analyses form the basis for the distance measures presented in the following section.

### 2.3 Two New Distance Measures on *m*-Clonal Trees

In this section, we first present some useful notation and then describe two novel tumor evolution distance measures restricted to *m*-clonal trees. In Section 2.4, we extend these measures to clonal trees in general.

#### 2.3.1 Notation

Suppose *T* ∈ ℑ is a clonal tree. We will need to translate between mutations and the vertices in a clonal tree that are labeled with those mutations. Therefore, given a mutation *i ∈ M*(*T*), we will denote the vertex in *T* ∈ ℑhat is labeled with mutation *i* as *v*_*i*_. We are interested in sets of mutations that label certain paths in a clonal tree. In particular, given *i ∈ M*(*T*), we define the *ancestral set A*(*i*) as the set of mutations that label the path from the root vertex *r* to *v*_*i*_ in *T*. This definition means that *A*(*i*) gives the set of mutations that exist in the clone represented by vertex *v*_*i*_. Given *i, j* ∈ *M* (*T*), we define the *common ancestor set C*(*i, j*) to be *A*(*i*) ∩ *A*(*j*). That is, *C*(*i, j*) is the set of mutations that are ancestral to both mutations *i* and *j*. Given *i, j* ∈ *M* (*T*_*k*_), we also define the *distinctly inherited set D*(*i, j*) to be *A*(*i*)\*A*(*j*). That is, *D*(*i, j*) is the set of mutations that are ancestral to mutation *i* but not mutation *j* in *T*. Note that under this definition it is almost always the case that *D*(*i, j*) ≠ *D*(*j, i*). When we have more than one tree, we use subscripts to distinguish between them. For instance, *A*_*k*_(*i*)*, C_k_*(*i, j*), and *D*_*k*_(*i, j*) all refer to *T*_*k*_. See Figure 1 for examples of ancestral sets, common ancestor sets, and distinctly inherited sets of a tree *T*_1_. Given two sets of mutations *A* and *B*, we note that the *Jaccard distance* between them is defined as 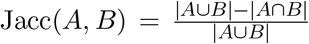 and Jacc(∅,∅) = 0.

**Figure 1:**
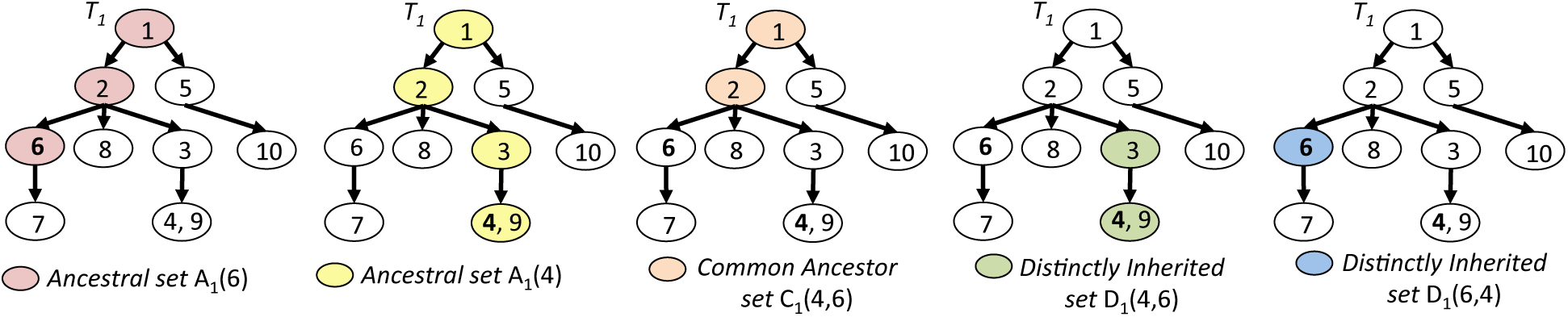
Examples of ancestral sets, common ancestor sets and distinctly inherited sets on one tree.

#### 2.3.2 Common Ancestor Set Distance

Given two *m*-clonal trees *T*_*k*_, *T*_*ℓ*_ ∈ ℑ_*m*_, we define a new tumor evolutionary tree distance measure called ***C**ommon **A**ncestor **Set** distance* (CASet) that computes a distance between *T*_*k*_ and *T*_*ℓ*_. Informally, CASet distance is the average Jaccard distance between all corresponding common ancestor sets in *T*_*k*_ and *T_ℓ_*. Equation (1) gives a formal definition of CASet distance.

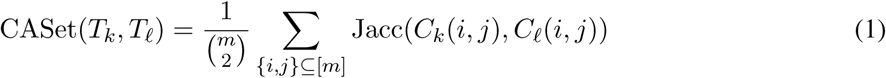

##### Observation 2.1.

*The running time to compute* CASet(*T*_*k*_, *T*_*ℓ*_) *is O*(*m*^3^).

A proof of Observation 2.1 can be found in the appendix. See Figure 2 for an example of CASet distance.

**Figure 2:**
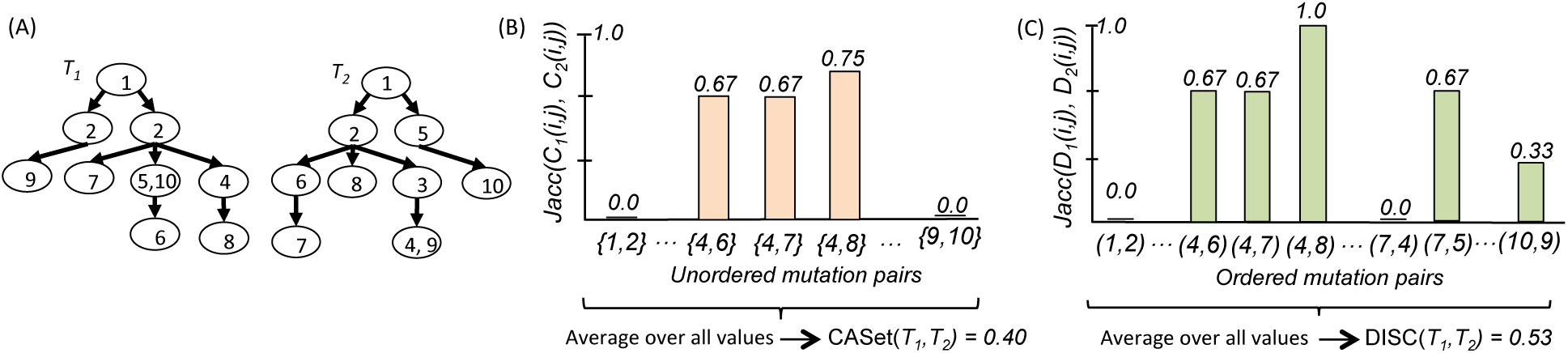
(A) A pair of 10-clonal trees. (B) Example of CASet distance applied to the 10-clonal trees in part (A). (C) Example of DISC distance applied to the 10-clonal trees in (A).

#### 2.3.3 Distinctly Inherited Set Comparison

CASet distance compares the common ancestor sets of all pairs of mutations, which emphasizes differences close to the root. However, we might also want to emphasize differences in more recently acquired mutations. Given two *m*-clonal trees *T*_*k*_, *T*_*ℓ*_ ∈ ℑ_*m*_, we define a new tumor evolution distance measure between *T*_*k*_ and *T*_*ℓ*_ called ***D**istinctly **I**nherited **S**et **C**omparison distance* (DISC) that accounts for mutation differences in the more recent tumor clones. Informally, DISC distance is the average Jaccard distance between all corresponding distinctly inherited ancestor sets in *T*_*k*_ and *T*_*ℓ*_. Equation (2) gives a formal definition of the DISC distance.

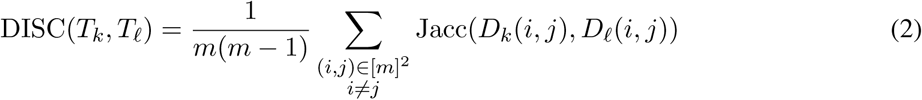

Note that this range of summation is different from CASet, which only considers unordered pairs {*i, j*}.

##### Observation 2.2.

*The running time to compute* DISC(*T*_*k*_, *T*_*ℓ*_) *is O*(*m*^3^).

The proof of Observation 2.2 is similar to the CASet runtime proof and can also be found in the appendix. Figure 2 also contains an example DISC computation.

#### 2.3.4 Algorithm Implementations

It is straightforward to implement these distance measures using set data structures to store all ancestor sets in the two trees. However, under certain circumstances, it may be faster or more convenient to compute CASet and DISC distances using matrix operations instead, which is described in Appendix A.2.

### 2.4 Extending CASet and DISC to Clonal Trees

Thus far, we have assumed that any two tumor evolutionary trees to be compared have the exact same set of mutation labels (that is, that they are both *m*-clonal). However, there are many scenarios in which this may not be the case. For instance, some methods such as [2] may not use all available mutations when creating a tumor evolutionary history tree. Further, different trees may be reconstructed from different data types for the same tumor (e.g. single cell and bulk sequencing in [31]) that do not share the same set of observed mutations. In this section, we present two extensions to both of our distance measures that allow for the comparison of clonal trees with different sets of mutation labels.

#### 2.4.1 Intersection of Mutation Sets

In the first extension to clonal trees, we consider the intersection of the mutation sets for the input trees. This allows us to compute a distance between two trees by only considering pairs of mutations that the two trees share. Let *I_k,ℓ_* = *M* (*T*_*k*_) ∩ *M*(*T*_*ℓ*_) be the intersection of the sets of mutations labeling *T*_*k*_ and *T*_*ℓ*_. Thus, we can modify both CASet and DISC distances as follows:

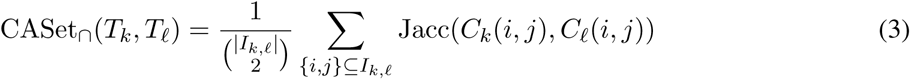

and

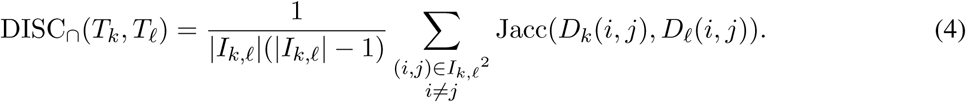

We note that while we restrict attention to pairs of mutations that exist in both trees, the actual sets compared using the Jaccard distance may themselves contain mutations that exist in only one of the trees. Thus, these distances are not the same as removing all non-shared mutations from the trees, contracting the tree topology, and computing the original version of the distances.

This variation is most useful when differences in mutation labelings between trees should not strongly contribute to their distance value. For example, this method could be used when comparing trees reconstructed using two different methods, one of which does not use all input mutations. A degenerate case arises if this approach is applied to two trees with disjoint mutation sets. Since we sum over pairs of shared mutations, CASet_∩_ and DISC_∩_ are both 0 in this case. If we want mutation set differences to be penalized, we can instead sum over pairs of mutations in the union of the two trees’ mutation sets, as described in the following section.

#### 2.4.2 Union of Mutation Sets

In the second extension to clonal trees, we consider the union of the mutation sets for the input trees. To do so, we need to address how to handle mutations that exist in only one tree. Let *U*_*k,ℓ*_ = *M* (*T*_*k*_) ∪ *M* (*T*_*ℓ*_) be the union of the sets of mutations labeling *T*_*k*_ and *T*_*ℓ*_. If *i* ∉ *M* (*T*_*k*_), then we define *A*_*k*_(*i*) = ∅ and compute *C*_*k*_(*i, j*) and *D*_*k*_(*i, j*) as usual. Thus, we can modify both CASet and DISC distances as follows:

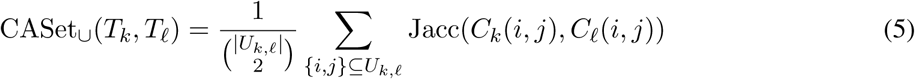

and

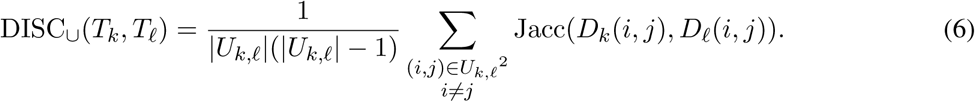

This variation allows differences in the sets of mutation labels to contribute to the distance computed between two trees. Thus, this variation may be most useful for comparing tumor evolutionary trees generated by different data types, across samples taken at different times, or even across patients. Notice that because Jacc(*X, ∅*) = 1 if *X* ≠ *∅*, we no longer have distance 0 between trees with disjoint labels. In the appendix, we describe a formula that relates CASet_∪_ and CASet_∩_, allowing us to compute CASet_*∪*_ with fewer operations.

## 3 Results

We compare CASet and DISC to existing distance measures on two simulated datasets and two real datasets. We find that our methods allow for more granularity in comparing tumor evolutionary trees and outperform other methods when used in a clustering context.

### 3.1 Results on Simulated Datasets

#### 3.1.1 Dataset Generation

We created two different simulated datasets for our analysis. In the first dataset, we manually constructed five “base” clonal trees each with 15 mutations but different topologies and labelings (Figure A.1), and generated five variants of each base tree for a total of 25 trees. The second dataset was generated using the OncoLib [32] tree generation tool, which simulates tumor evolutionary trees and read count data. We took the read count data from five OncoLib simulations (each with 3 sequenced samples and 10–20 mutations) and reconstructed clonal trees that were consistent with the simulated data [22]. Each OncoLib simulation yielded a “tree family” of between 50 and 10,000 clonal trees, of which 50 were sampled to create a dataset of 250 trees from five families (labeled A–E). All trees within a family have the same set of mutations, but mutation sets differ across families. The true underlying clonal trees produced by OncoLib are in Figure A.5.

#### 3.1.2 Correlation Between CASet and DISC

In each of the five tree families from the OncoLib dataset, the underlying tree structure has a strong effect on the correlation between CASet and DISC distances (Figure 3). In general, two trees with a large CASet distance also have a large DISC distance; however, family A includes some pairs of trees whose CASet and DISC distances differ significantly.

**Figure 3:**
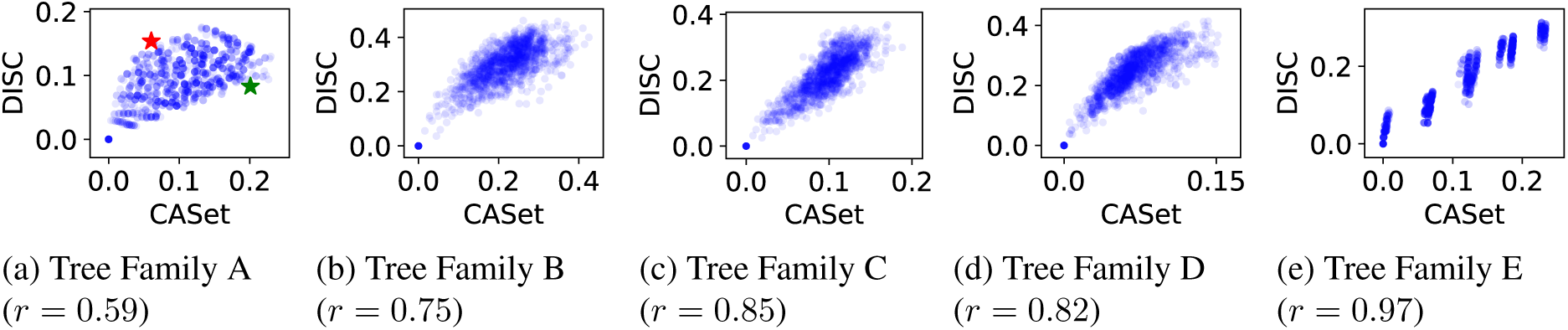
Correlation of CASet and DISC on tree families A-E on the OncoLib dataset. Pearson correlation coefficients are shown in parentheses. The red and green stars in (a) mark the tree pairs *T*_14_, *T*_26_ and *T*_5_, *T*_20_, respectively. Note that *x* and *y* scales are not consistent across all plots in order for all patterns to be visible.

The different patterns of CASet and DISC correlation are related to the height of the underlying true tree. For example, the true tree underlying family A (Figure A.5) is the shallowest of the set and has the lowest correlation between CASet and DISC, while the true tree of family E is the deepest tree and has the highest correlation. Shallow trees will tend to be wider, and therefore the ancestral set comparisons used by CASet will frequently only contain the root (whose labels are consistent due to the tree reconstruction process). As such, the Jaccard distance between the two sets will be 0 more frequently than in a deeper tree, in which more labels are in the ancestral set on average, causing any given label change to have an impact across more sets. In contrast, changes in shallow trees (with many leaves) will be weighted slightly more heavily by DISC by a similar argument. The relationship between tree height and distance correlation can also be seen for the manually constructed dataset (Figure A.2).

We then examined two pairs of trees in the OncoLib dataset A for which CASet and DISC disagreed significantly, *T*_14_, *T*_26_ and *T*_5_, *T*_20_, highlighted in Figure 3a. According to CASet, *T*_14_ and *T*_26_ are more similar than *T*_5_ and *T*_20_, while DISC reports the opposite (Figure 4). The CASet distance between *T*_14_ and *T*_26_ is small, since ancestral relationships in the two trees are nearly identical (label 3 being the exception). However, the larger DISC distance between *T*_14_ and *T*_26_ reflects the significant differences in leaf locations in the two trees. Nine mutations are direct children of the root in both *T*_14_ and *T*_26_, but eight mutations are children of 3 in exclusively one tree. This results in a large number of mutation pairs with different sets of distinctly inherited ancestors, causing the DISC distance to be comparatively large. On the other hand, the CASet distance between *T*_5_ and *T*_20_ is large, reflecting the significant ancestral differences between the trees. Many more mutations have 3 as an ancestor in *T*_20_ than in *T*_5_. However, the DISC distance is relatively small because enough siblings move together between the two trees that their sets of distinct ancestors are mostly similar. Thus we see that CASet emphasizes ancestral similarity and performs especially well on deeper trees, while DISC prioritizes similarity in leaf relationships and is more granular with shallower trees.

**Figure 4:**
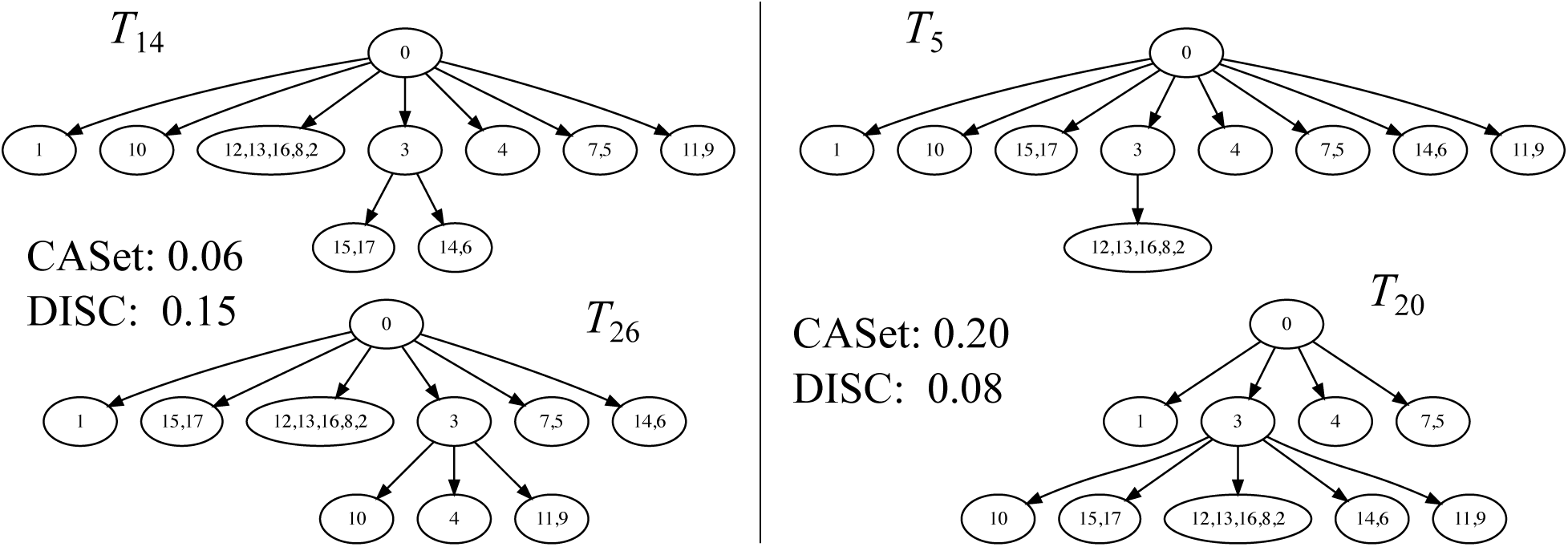
CASet and DISC calculations for trees *T*_14_ and *T*_26_ (left) and *T*_5_ and *T*_20_ (right) from the OncoLib dataset. These pairs of trees are starred in Figure 3(a).

#### 3.1.3 Clustering Clonal Trees

Previous work has shown that many different hypothesized mutation trees can be consistent with data from a single patient [21, 22]. Clustering these trees is a compelling use case for tumor evolution distance measures as it has the potential to reveal structure in the space of compatible trees of a single dataset. Clustering trees inferred from different patients can also be used to identify shared evolutionary patterns. We compare our distance measures to MLTED [28], the four distance measures introduced in [20] (ancestor-descendant, parentchild, clonal, and path distances), and triplet distance [33] (a modified version of a distance designed for phylogenetic trees, described in A.4) using a clustering scenario on both simulated datasets.

We performed hierarchical clustering with average linkage on both the manual and OncoLib datasets. On the manual dataset, we compute the average silhouette value [34] for different cuts of the resulting tree on all tested distance measures and use the cut with the highest such value to produce a clustering of the data (Figure 5). Heatmaps showing all pairwise distances are in Figure A.4. We found that several of the methods (including CASet and DISC) produce the correct number of clusters, five.

**Figure 5:**
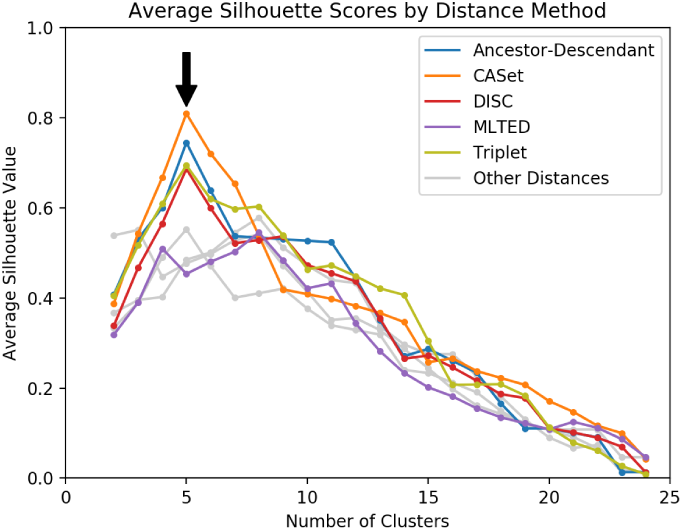
Plot of average silhouette values for different cuts of the hierarchical clustering with average linkage tree for each distance method on the manually created dataset. Three distances (clonal, parentchild, and path) did not have a large silhouette score for 5 clusters and are shown in gray. See Figure A.3 for a fully labeled plot.

On the OncoLib dataset, we applied CASet_∪_, CASet_∩_, DISC_*∪*_, DISC_*∩*_, and MLTED [28] when doing hierarchical clustering. Figure 6 shows heatmaps of all pairwise tree distances for the five tree families A–E and the average silhouette score for five clusters, which all but one method identified as the optimal hierarchical clustering cut (CASet_∩_ had an optimal cut at three clusters). When cut at five clusters, all five distance measures correctly clustered the trees, but they did so with varying degrees of tightness. CASet_∪_ performed best in distinguishing trees belonging to different datasets, with an average silhouette score of 0.81 over the five tree families. While it performs worse at separating different families, CASet_∩_ identifies that the pairs A, B and D, E have strong agreement about ancestral relationships among their shared mutations (see Figure A.5). This highlights the different useful features of CASet_∪_ and CASet_∩_. In comparison, MLTED did a worse job of separating trees from different groups, with an average silhouette score of 0.54 over the five tree families. At the same time, it also does not recognize that the relationships between mutations shared by the D and E tree families are very similar.

**Figure 6:**
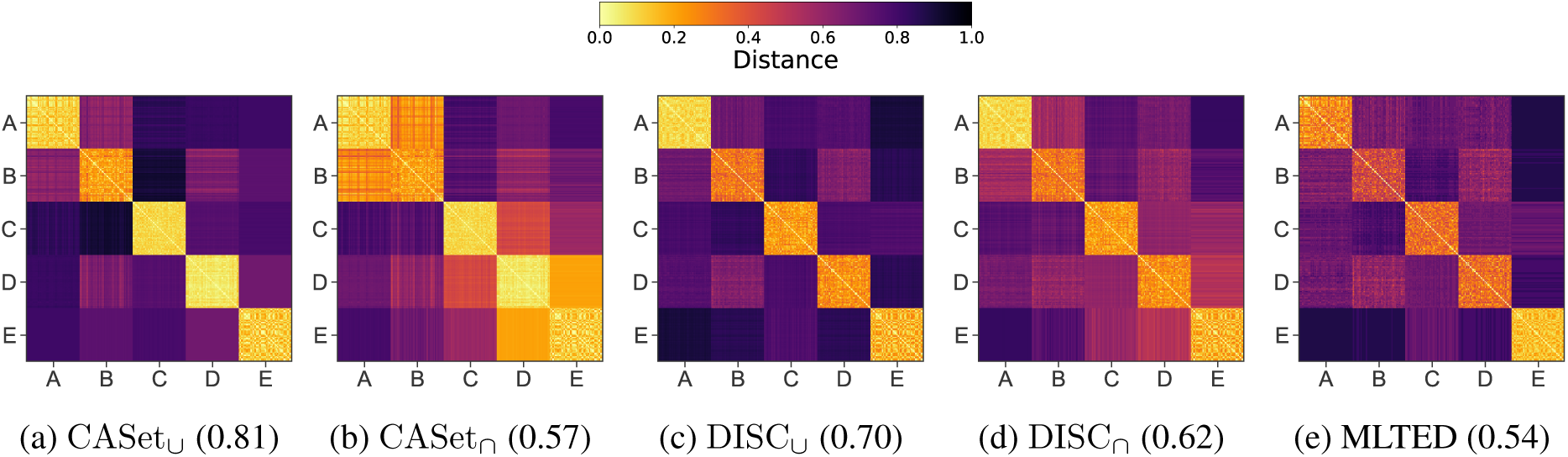
Inter-dataset distance heatmaps of five tree families in the OncoLib dataset. The color of each cell represents the distance between two trees. Average silhouette scores for five clusters are displayed in parentheses to quantify clustering tightness and separation.

#### 3.1.4 Intra-family Clustering Structure

Figures 3e and 6 show that tree family E from the OncoLib dataset might have internal structure. Therefore we took a closer look at the internal clustering structure for this tree family (see Figure 7). In this dataset, CASet, DISC, and MLTED all identify two primary clusters of trees, but CASet and DISC are both able to resolve more complex substructure. In particular, CASet distinguishes between eight strongly defined subfamilies, with an average silhouette score of 0.94. The same eight subfamilies are visible in the DISC heatmap along with a finer-grained resolution within each of these clusters. In contrast, the MLTED heatmap shows less resolution within the two primary categories. For a similar set of plots of family A, see Figure A.7, but none of the other four tree families had internal structure as well-defined as family E under any of the three distances.

**Figure 7:**
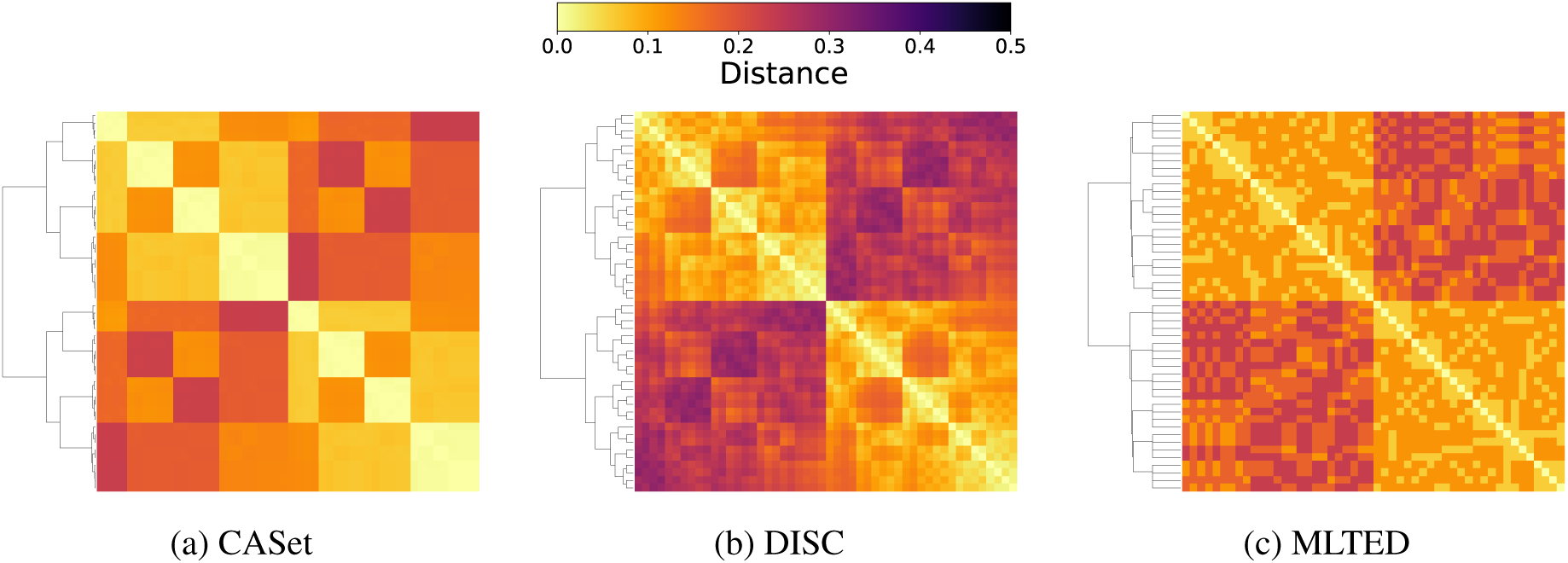
Clustering of dataset E. Note that the colormap range has been reduced to provide more contrast.

### 3.2 Results on Real Datasets

We apply our distance measures to two different breast cancer datasets [36, 37]. We first apply CASet and DISC to the three potential tumor evolutionary histories reported in [28] recovered using different methods applied to single cell sequencing data from a triple negative breast cancer patient [36] (see Figure 8). Since CASet and DISC are designed to evaluate the topology of trees in addition to their labels and inheritance, both measures are able to provide more granular information about the similarity of the trees making it possible to conclude that *T*_2_ is more similar to *T*_1_ than it is to *T*_3_. This is in contrast to MLTED [28], which considers these pairs of trees to have the same similarity, and, furthermore, computes the distance between *T*_1_ and *T*_3_ as 0 since they can be clonally expanded to match. While it may be useful to evaluate the similarity of trees with regard to such clonal expansion, a tumor evolutionary tree is categorized in part by subclonal populations that represent the evolutionary patterns of the tumor. Ignoring these does not fully take into account the information represented in a tumor evolutionary tree.

**Figure 8:**
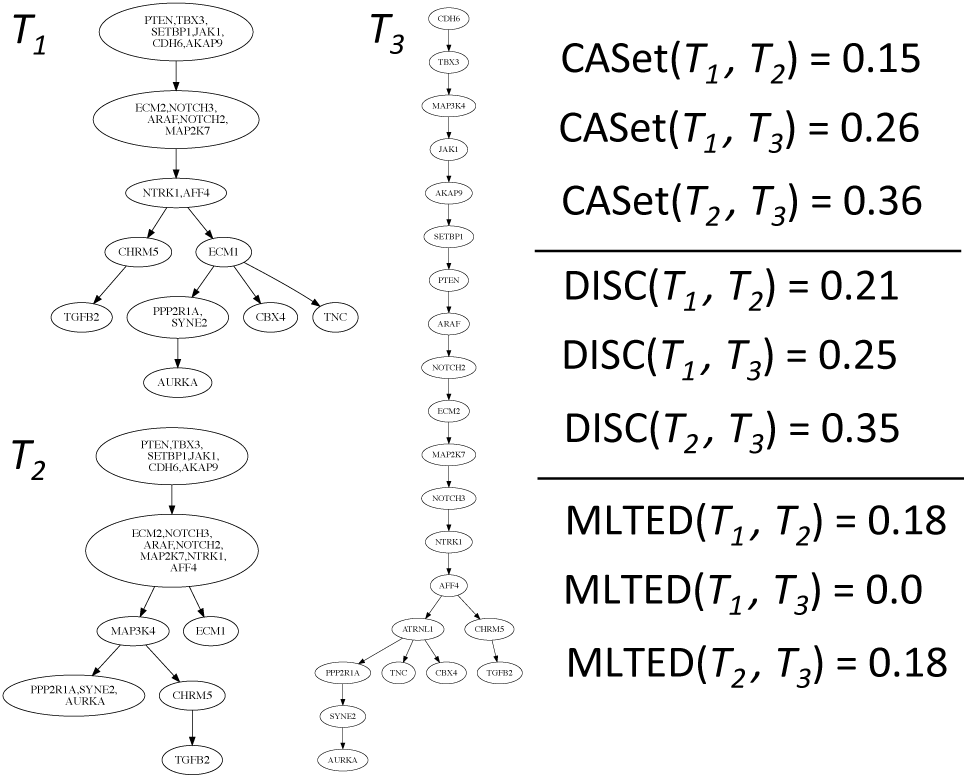
Tumor evolutionary trees inferred by PHiSCS [35] (*T*_1_), SciFit [13] (*T*_2_), and SCITE [10] (*T*_3_) from a triple negative breast cancer patient [36] as reported by [28] and corresponding pairwise distances.

We also emphasize the importance of using quantitative measures when comparing reconstructed trees. The authors of [8] introduced a new tree reconstruction method MIPUP, and when they compare their results to those of another method, LICHeE [3], on breast cancer xenoengraftment data sample SA501 [37], they use only qualitative analysis to claim their method produces phylogenies closer to those in the original publication. We assessed this claim quantitatively by running CASet, DISC, and MLTED on the SA501 tree from [37] and the corresponding trees reconstructed by MIPUP and LICHeE, as shown in [8]. These trees had around 180 SNV mutations each in 5-7 nodes with slightly different mutation sets between trees. We found that the LICHeE tree was more similar to the phylogeny proposed by [37] than the MIPUP tree according to both CASet and DISC. The CASet/DISC distances were 0.837/0.397 for the MIPUP tree and 0.781/0.377 for the LICHeE tree. However, MLTED reports the opposite by a small margin, with distances of 0.798 for the MIPUP tree and 0.811 for the LICHeE tree compared to the proposed phylogeny. We note that with trees of this size, distances of this magnitude are more common than of the magnitudes represented in the OncoLib dataset, for example. However, the purpose of this analysis is to determine the relative distances of two trees to a ground truth tree and the raw distance is less relevant. Therefore, although the MIPUP tree may be marginally more similar to the originally proposed tree from a tree edit distance standpoint, it appears to be less similar when we account for the ancestral patterns of the disagreements.

## 4 Discussion and Conclusion

In this work, we argue that distance measures designed for tumor evolutionary trees are needed for assessing phylogeny inference methods and for exploring the relationships between sets of evolutionary trees. To this end, we introduce two new tumor evolution distance measures, CASet and DISC. By comparing the common ancestors of all mutation pairs, CASet incorporates differences in both mutation labeling and tree topology. In particular, CASet uses the number of clones that inherit common mutations when weighting the effect of mutation labeling differences between trees. In contrast, DISC pays special attention to the set of mutations that distinguish clones from each other, placing comparatively more emphasis on recently acquired mutations. We extend both distance measures to apply clonal trees with different sets of mutations.

We demonstrate the differences between CASet and DISC on simulated data and use a clustering application to show that that CASet is better able to distinguish groups of trees than existing distance measures. Moreover, we find that both CASet and DISC can identify complex clustering structure in a space of trees that is missed by the MLTED distance measure [28].

Using a breast cancer dataset, we show that CASet and DISC are able to differentiate between trees with differing clonal makeups, demonstrating their topological acuity. In addition, we use CASet and DISC to assess trees reconstructed by MIPUP [8] and LICHeE [3] from a breast cancer xenograft dataset [37]. Our results suggest that the MIPUP tree may not more closely resemble the original hypothesized tree from [37] than the LICHeE tree, as is suggested in [8] based solely on qualitative analysis. This highlights the importance of using quantitative measures such as CASet and DISC for these types of assessments.

Future work is needed to determine the benefits of using CASet or DISC in methods like GraPhyC [20] that rely explicitly on distance measures. Our findings on the internal clustering structure of the OncoLib tree families also invite further investigation. It remains to be seen whether the same kind of structure is found in real cancer data—either within sets of trees consistent with data from a single patient or across sets of patients with the same type of cancer. If so, this structure could provide insight into improved tumor phylogeny inference methods or common mutational patterns across patients.

## A Appendix

## A.1 Runtime Proofs

*Proof of Observation 2.1.* We first do some pre-processing by computing *A*_*k*_(*i*) for all *i* ∈ [*m*] using a depth first traversal of *T*_*k*_. We keep track of the set of mutations on the path from the root to the current vertex while executing the algorithm. When visiting vertex *v* during the traversal, we can then store *A*_*k*_(*i*) for all mutations *i* that label vertex *v*. Since we store *m* sets of mutations, each of which may contain up to *m* mutations, this step takes *O*(*m*^2^). Next we sort each of these *m* sets which takes *O*(*m*^2^ log *m*). Next we can compute the 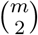 common ancestor sets, each of which takes *O*(*m*) to find since the Ancestor sets are sorted. This step is then *O*(*m*^3^). We do the same steps for *T*_*ℓ*_. Finally we need to compute the Jaccard distance between each of the 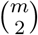 common ancestor sets for *T*_*k*_ and *T*_*ℓ*_. Since all these sets are in sorted order, each of these computations is *O*(*m*). Thus, computing these Jaccard distances and summing them up to get the CASet distance is *O*(*m*^3^). Therefore, the overall runtime to compute CASet distance is *O*(*m*^3^).

*Proof of Observation 2.2.* We first do some pre-processing by computing *A*_*k*_(*i*) for all *i* ∈ [*m*] by doing a depth first traversal of *T*_*k*_. We keep track of the set of mutations on the path from the root to the current vertex while executing the algorithm. When visiting vertex *v* during the traversal, we can then store *A*_*k*_(*i*) for all mutations *i* that label vertex *v*. Since we store *m* sets of mutations, each of which may contain up to *m* mutations, this step takes *O*(*m*^2^). Next we sort each of these *m* sets which takes *O*(*m*^2^ log *m*). Next we can compute the *m*^2^ distinctly inherited sets, each of which takes *O*(*m*) to find since the Ancestor sets are sorted. This step is then *O*(*m*^3^). We do the same steps for *T*_*ℓ*_. Finally we need to compute the Jaccard distance between each of the *m*^2^ distinctly inherited ancestor sets for *T*_*k*_ and *T*_*ℓ*_. Since all these sets are in sorted order, each of these computations is *O*(*m*). Thus, computing these Jaccard distances and summing them up to get the DISC distance is *O*(*m*^3^). Therefore, the overall runtime to compute DISC distance is *O*(*m*^3^).

## A.2 Matrix Implementation

Let *A*_*k*_ denote the *m × m* binary *ancestor matrix* of *T*_*k*_, which contains a 1 in entry *A*_*k*_[*i, j*] if mutation *i* is an ancestor of mutation *j*. To compute CASet distance, we first have to find the common ancestors of every pair of columns in *A*_*k*_ and in *A*_*ℓ*_. This can be done quickly by taking boolean ANDs of columns. Similarly, distinctly inherited sets can be found using XORs of columns. Then, the Jaccard distances in the algorithm can be computed by performing boolean ANDs and ORs on the common ancestor/distinctly inherited vectors to find intersections and unions, respectively, and finally summing the entries. Note that the process of computing all common ancestors within a tree should be performed as a pre-processing step if a large number of trees are being compared to each other. Lastly, we note that the asymptotic running time of this approach is *O*(*m*^3^), but it may still be faster in practice depending on available hardware and programming language specifics.

## A.3 A Direct Relationship Between CASet_∪_ and CASet_∩_

In Theorem 1, we describe a direct relationship that allows us to compute CASet_∪_ based on CASet_∩_. In particular, CASet_∪_ is a weighted sum of CASet_∩_ with terms that quantify the number and proportion of mutations unique to each tree. Using Theorem 1, we can compute CASet_∩_(*T*_*k*_, *T*_*ℓ*_) and then use the formula to compute CASet_∪_(*T*_*k*_, *T*_*ℓ*_) if desired. This allows us to consider fewer pairs *i, j*, which may be helpful in practice when the trees have significantly different sets of mutations. The relationships between mutations considered by DISC are more complex, which prevents us from performing a similarly clean expansion of DISC_*∪*_.

### Theorem 1.

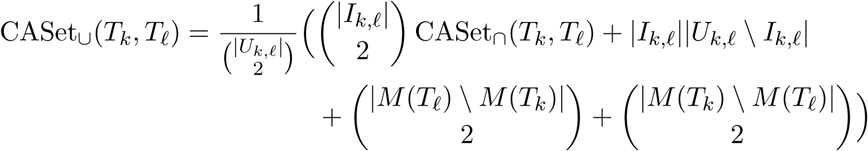

*Proof of Theorem 1.* By definition,

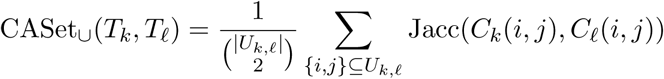

We can break this sum into parts (a)-(d), based on the cases for *i* and *j*:

a. *i, j* ∈ *I*_*k,ℓ*_
b. *i* ∈ *I*_*k,ℓ*_ and *j* ∈ *I*_*k,ℓ*_
c. *i, j* ∈ *M* (*T*_*ℓ*_) \ *M* (*T*_*k*_) or *i, j ∈ M* (*T*_*k*_) \ *M*(*T*_*ℓ*_)
d. *i* ∈ *M* (*T*_*ℓ*_) \ *M* (*T*_*k*_) and *j* ∈ *M* (*T*_*k*_) \ *M* (*T*_*ℓ*_)

Notice that if either *i* or *j* does not appear in *T*_*k*_, we will have *C*_*k*_(*i, j*) = ∅, and likewise for *T*_*ℓ*_. Thus, in cases (b) and (c), we have Jacc(*C*_*k*_(*i, j*), *C*_*ℓ*_(*i, j*)) = 1. However, in case (d), the Jaccard distance is Jacc(*∅*, *∅*) = 0. We’ll use these case labels to abbreviate our summations. Using these facts and the definition of CASet_*∩*_,

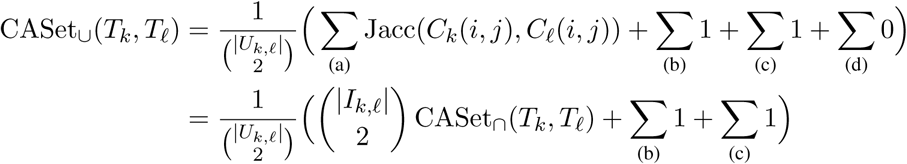

Finally, notice that the number of pairs *i, j* in case (b) is |*I*_*k,ℓ*_║*U*_*k,ℓ*_ \ I_*k,ℓ*_| and that the number of pairs in 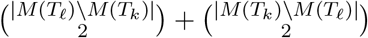. We can then explicitly write out the (b) and (c) sums:

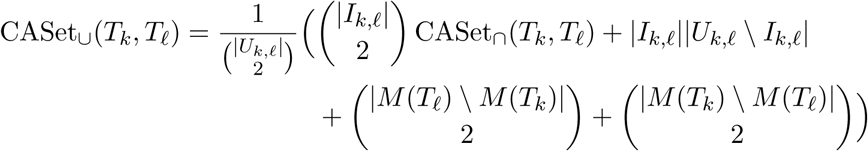

## A.4 Extending Triplet Distance to Tumor Evolutionary Trees

The triplet distance is an algorithm designed to compute the similarity of phylogenetic trees based on subtree relationships [27]. For every triplet of leaf nodes (in a phylogenetic tree, each leaf node has exactly one label), the closer two nodes (determined via the last common ancestor of each pair) are recorded. The count of triplets for which this closeness does not match across trees is normalized by the total number of comparisons made, 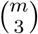 for an *m*-clonal tree, and recorded as the distance. It is possible to adapt this method to a tumor evolutionary tree by picking all triplets of mutation labels and ranking proximity based on the distance of their last common ancestors to the root.

## A.5 Additional Figures

**Figure A.1:**
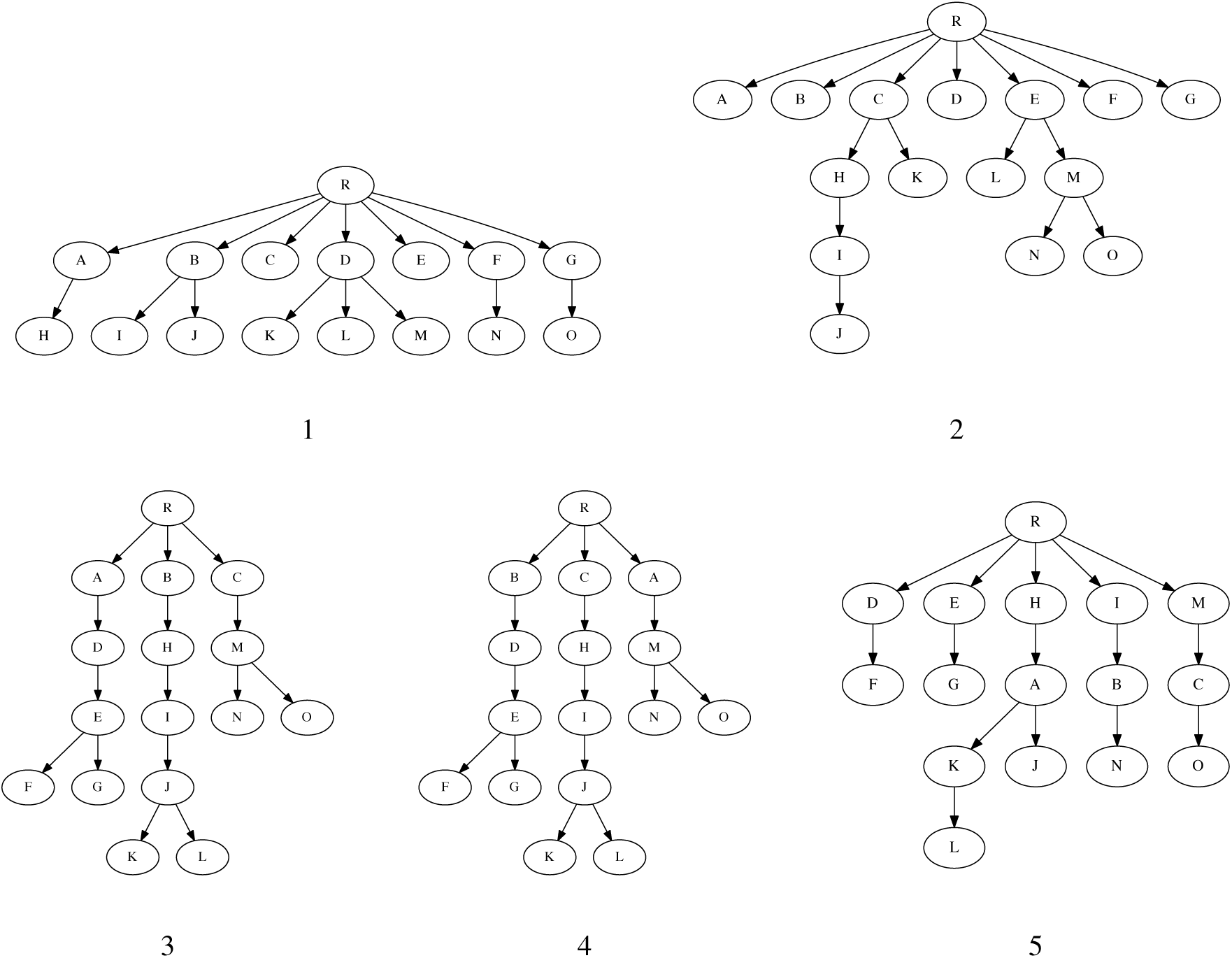
The base trees for the manual clustering experiment, in which five variants of each of these trees were created.

**Figure A.2:**
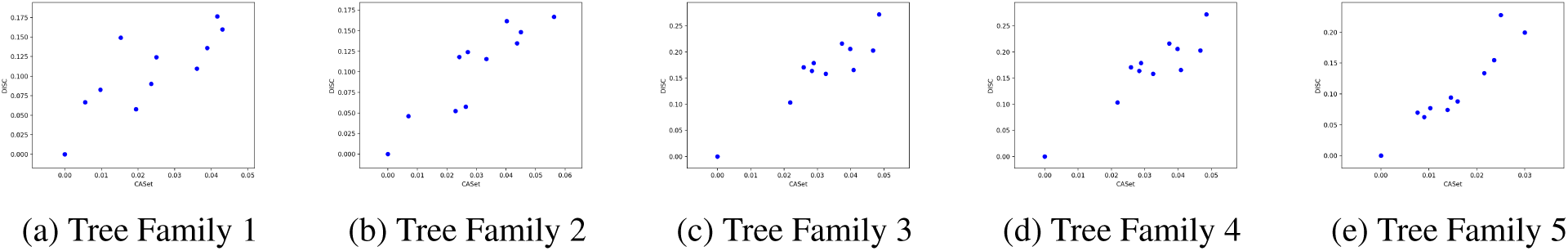
Scatterplots of CASet and DISC correlation on the manual dataset. Again, the shallower trees (families 1 and 2) have the greatest disparity between distance measures (see Section 3.1.2 for an elaboration).

**Figure A.3:**
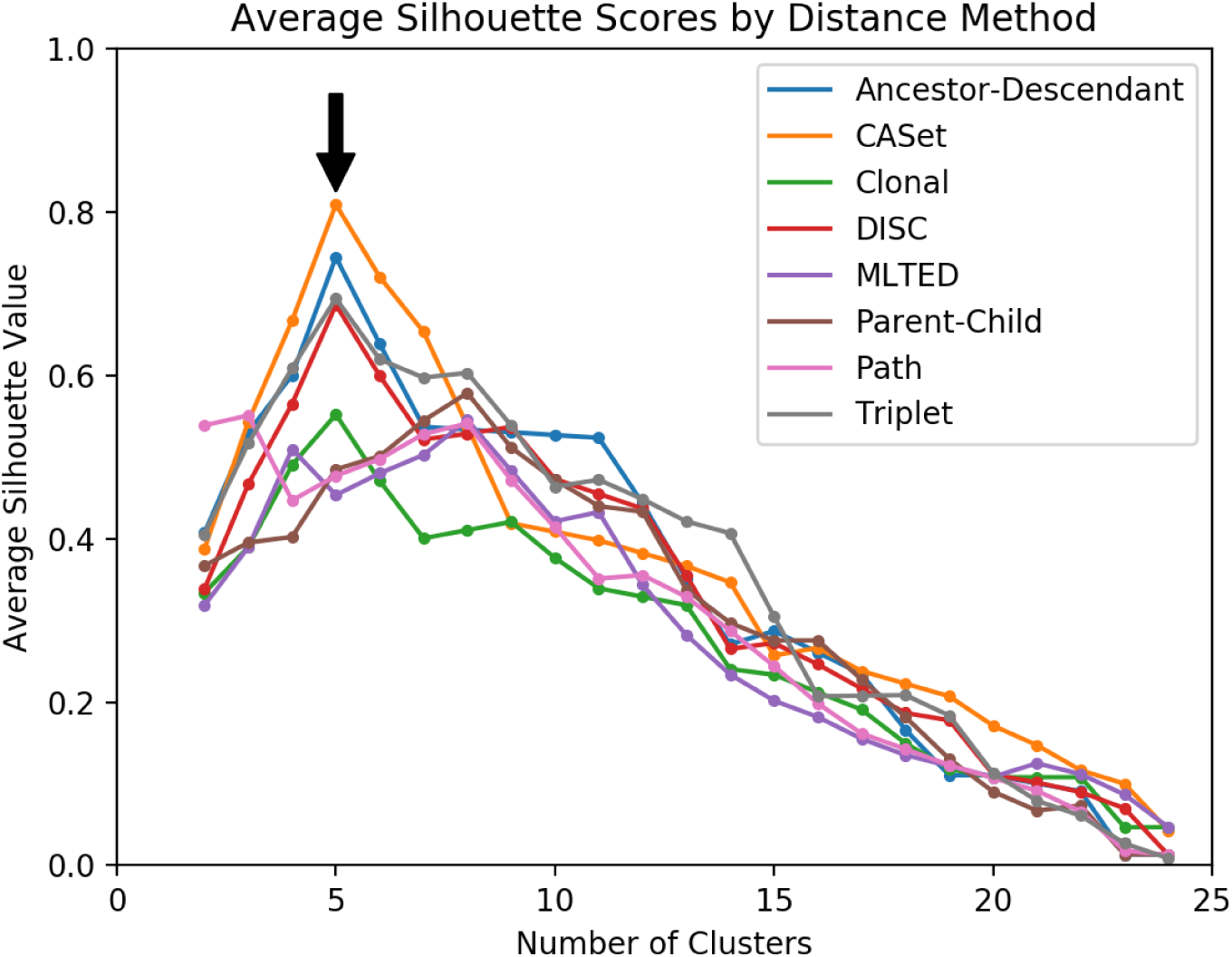
All silhouette plots for the manual dataset.

**Figure A.4:**
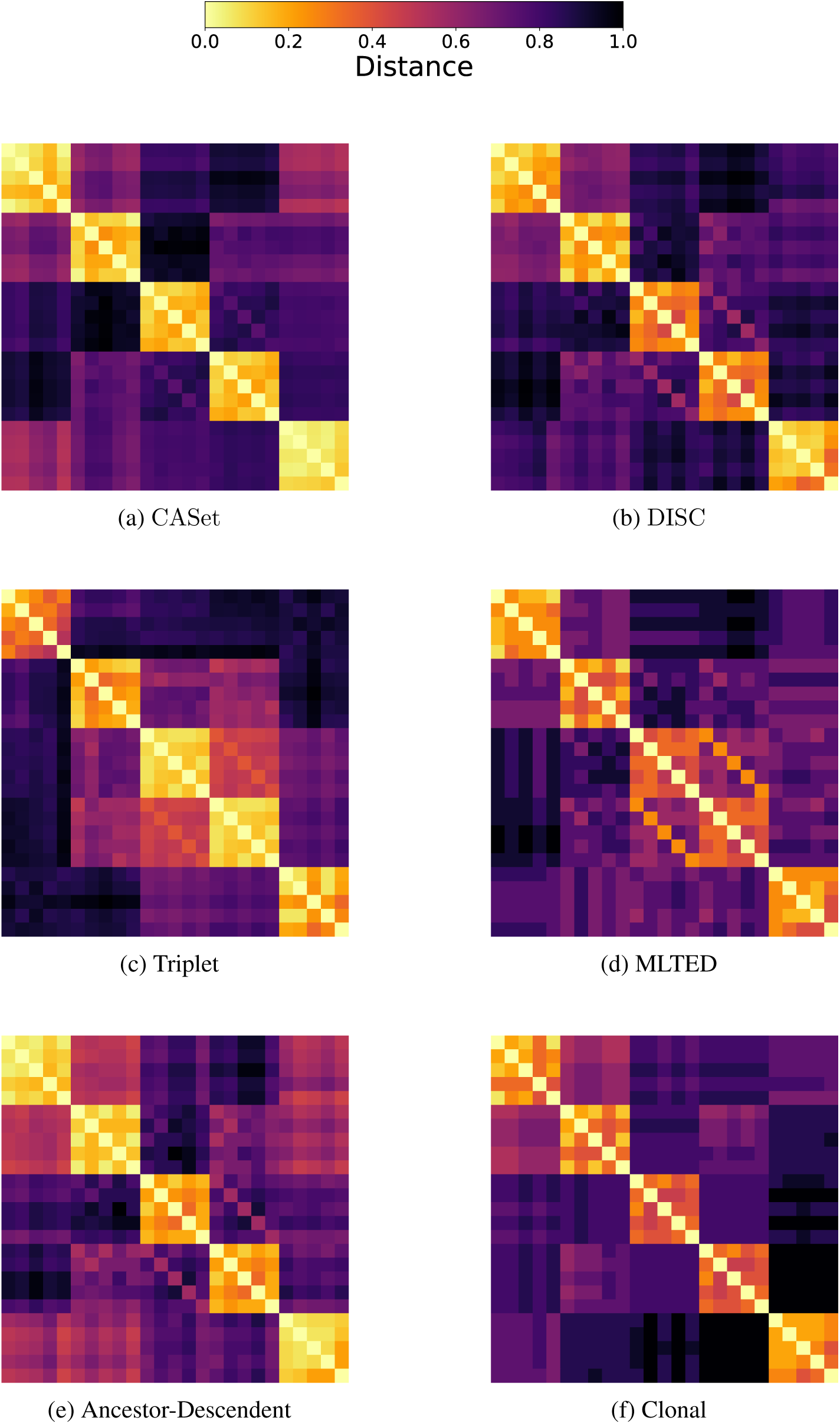
Heatmaps showing the (unclustered) pairwise distances between trees. In each, lighter shades represent closer trees. Defined 5-by-5 boxes show that the method measures trees within a cluster as being very similar to each other and less similar to other groupings.

**Figure A.5:**
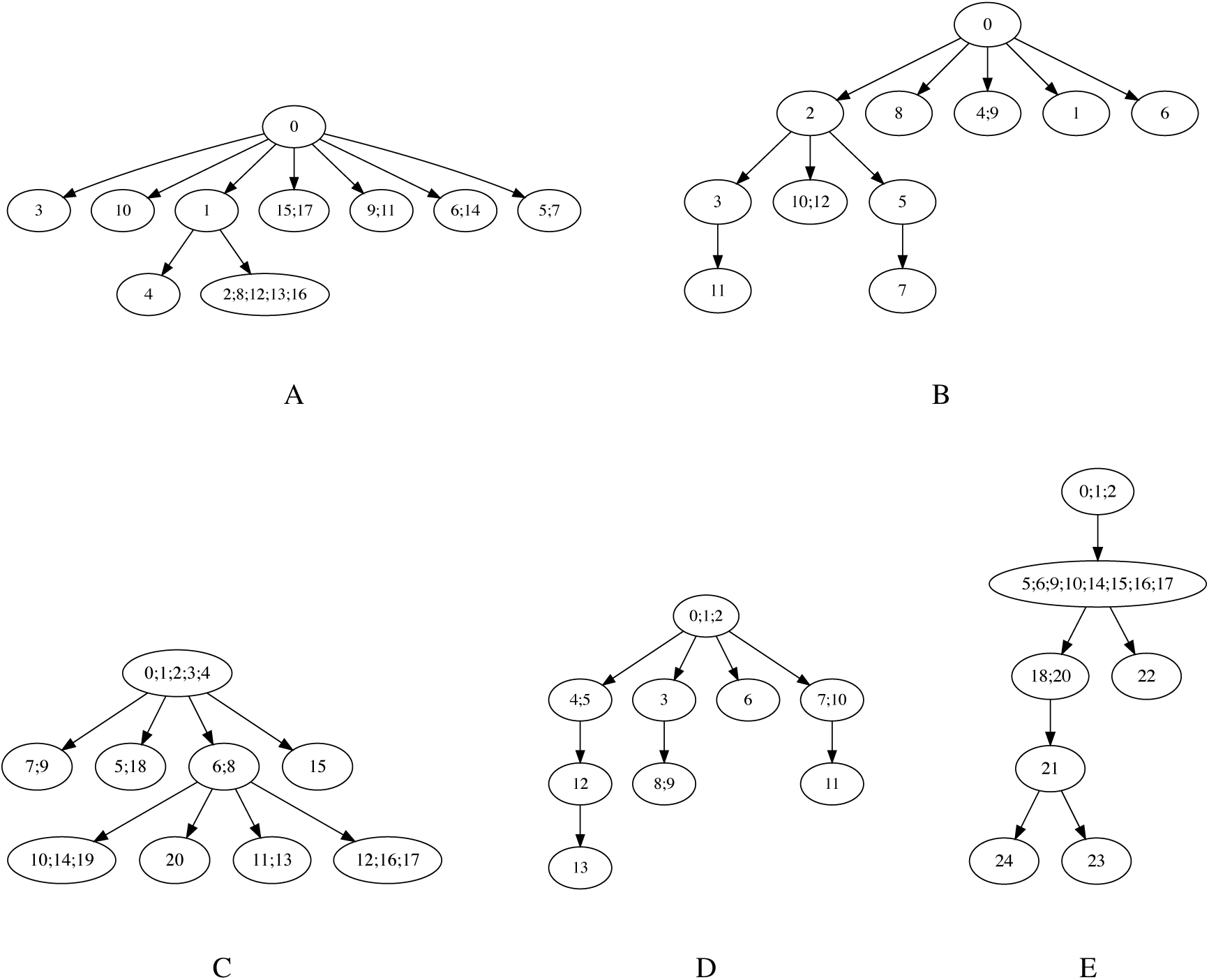
Mutation trees from OncoLib. OncoLib simulations are structured such that mutations are represented by ordered integers, so all five tree families had at least 10 common mutations, often near the root of the tree.

**Figure A.6:**
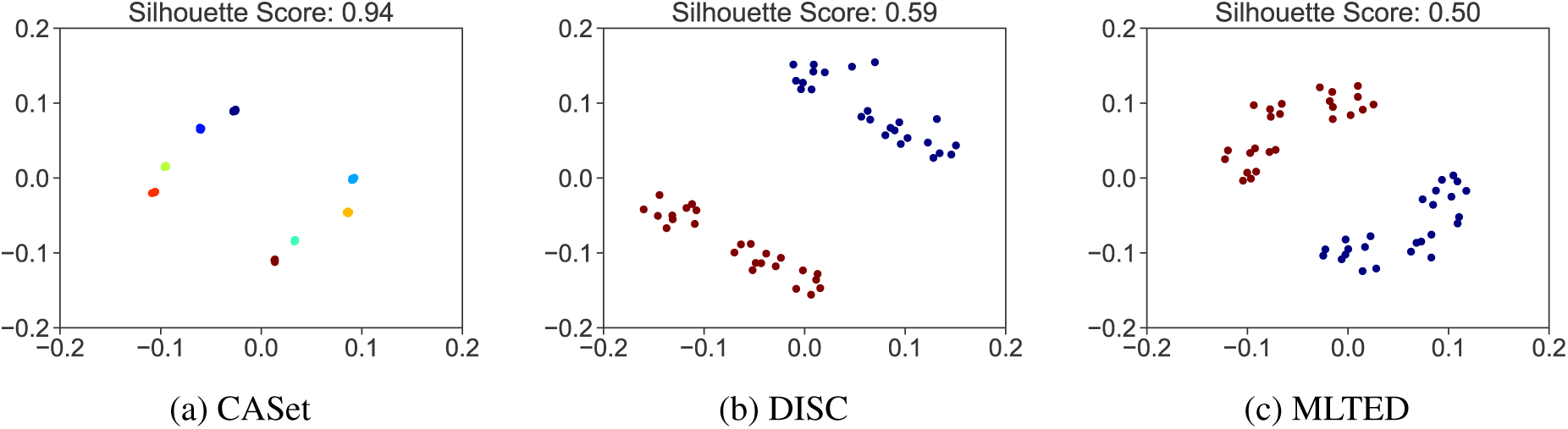
Multidimensional scaling (MDS) plots of OncoLib family E. These provide another way of visualizing the clustering in Figure 7. Colors correspond to clusters in the optimal silhouette cut, the score for which is shown above each plot.

**Figure A.7:**
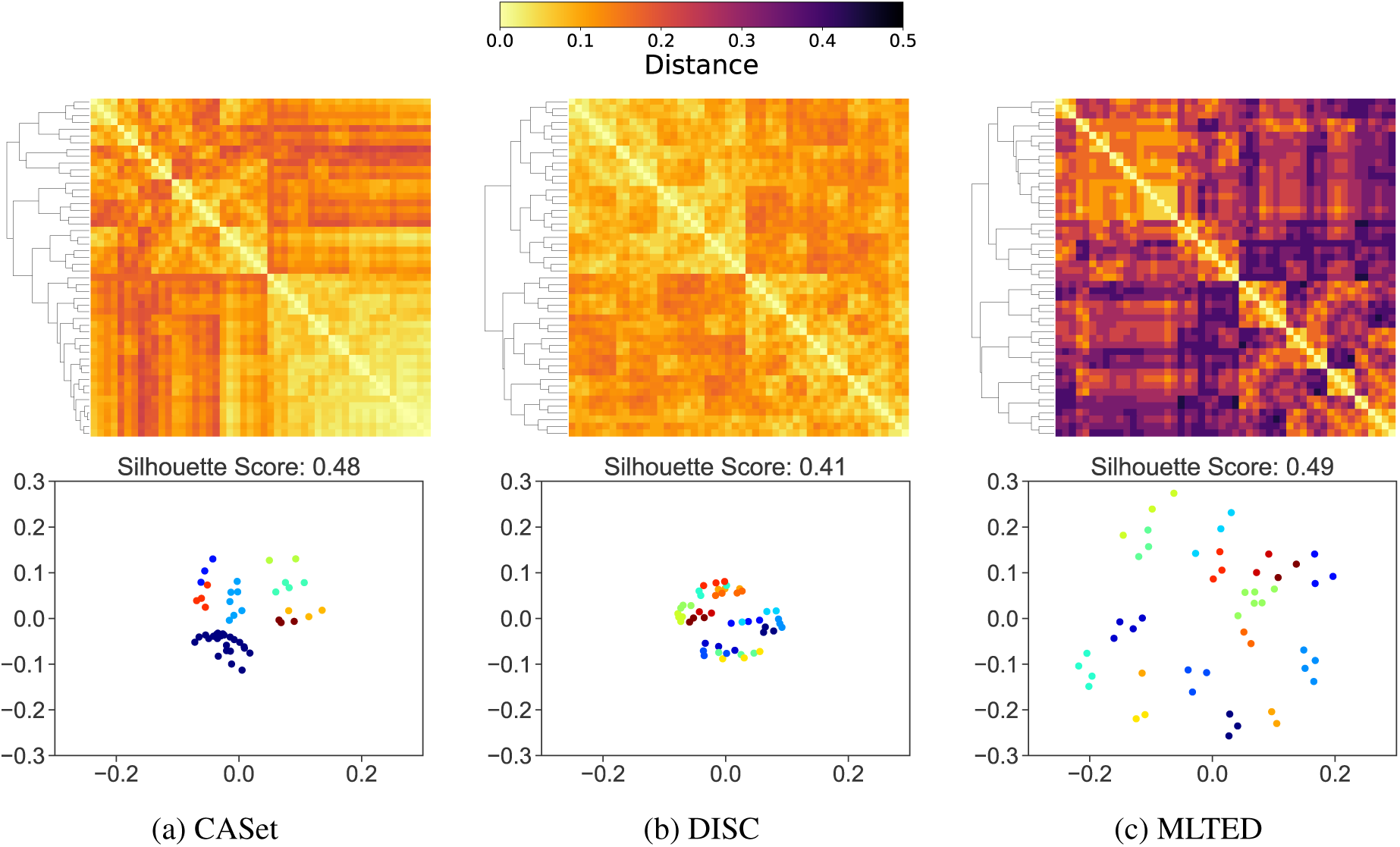
Clustering of dataset A. The top row shows hierarchically clustered distance heatmaps and the bottom row shows multidimensional scaling plots.

**Figure A.8:**
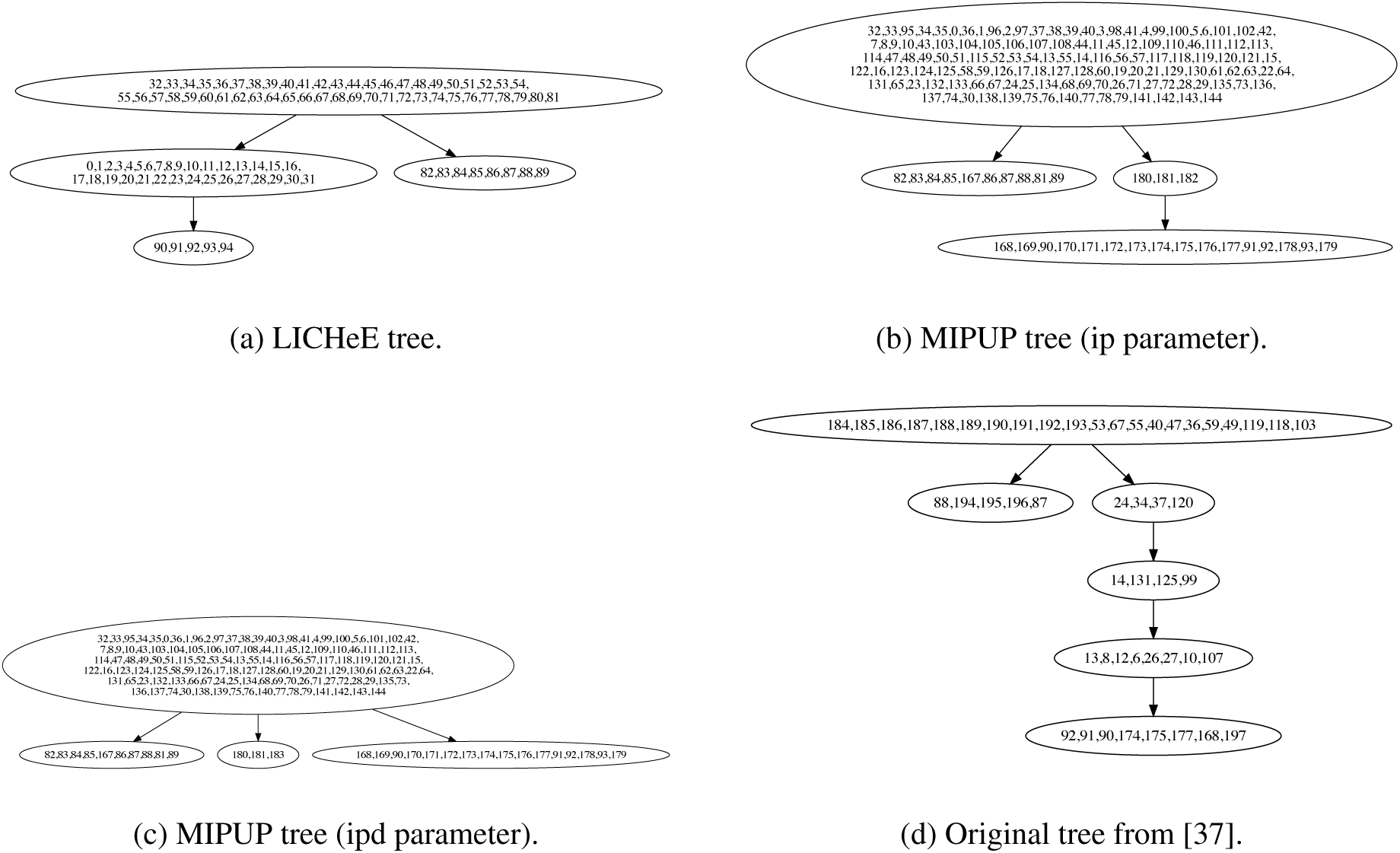
Trees inferred from the SA501 dataset presented in [8]. We used tree (b) when referring to the MIPUP tree. The distances to tree (c) differed by an insignificant amount from the distances we reported (less than 0.01).

## Notes

This project is supported by NSF award IIS-1657380, Elledge, Eugster, and Class of’49 Fellowships from Carleton College

## Reference

[1] P. C. Nowell. The clonal evolution of tumor cell populations. Science, 194(4260):23–8, Oct 1976.

[2] Mohammed El-Kebir, Layla Oesper, Hannah Acheson-Field, and Benjamin J. Raphael. Reconstruction of clonal trees and tumor composition from multi-sample sequencing data. Bioinformatics, 31(12):i62–i70, Jun. 2015.

[3] Victoria Popic, Raheleh Salari, Iman Hajirasouliha, Dorna Kashef-Haghighi, Robert B West, and Ser-afim Batzoglou. Fast and scalable inference of multi-sample cancer lineages. Genome Biol, 16:91, May 2015.

[4] Salem Malikic, Andrew W McPherson, Nilgun Donmez, and Cenk S Sahinalp. Clonality inference in multiple tumor samples using phylogeny. Bioinformatics, 31(9):1349–56, May 2015.

[5] Gryte Satas and Benjamin J Raphael. Tumor phylogeny inference using tree-constrained importance sampling. Bioinformatics, 33(14):i152–i160, Jul 2017.

[6] Wei Jiao, Shankar Vembu, Amit G Deshwar, Lincoln Stein, and Quaid Morris. Inferring clonal evolution of tumors from single nucleotide somatic mutations. BMC Bioinformatics, 15:35, Feb 2014.

[7] Hosein Toosi, Ali Moeini, and Iman Hajirasouliha. BAMSE: Bayesian model selection for tumor phylogeny inference among multiple tumor samples. In 2017 IEEE 7th International Conference on Computational Advances in Bio and Medical Sciences (ICCABS), pages 1–1. IEEE, 2017.

[8] Edin Husić, Xinyue Li, Ademir Hujdurović, Miika Mehine, Romeo Rizzi, Veli Mäkinen, Martin Milanič, and Alexandru I Tomescu. MIPUP: Minimum perfect unmixed phylogenies for multi-sampled tumors via branchings and ILP. Bioinformatics, 2018.

[9] Noushin Niknafs, Violeta Beleva-Guthrie, Daniel Q Naiman, and Rachel Karchin. Subclonal hierarchy inference from somatic mutations: Automatic reconstruction of cancer evolutionary trees from multi-region next generation sequencing. PLoS Comput Biol, 11(10):e1004416, Oct 2015.

[10] Katharina Jahn, Jack Kuipers, and Niko Beerenwinkel. Tree inference for single-cell data. Genome Biol, 17:86, May 2016.

[11] Sayaka Miura, Louise A Huuki, Tiffany Buturla, Tracy Vu, Karen Gomez, and Sudhir Kumar. Computational enhancement of single-cell sequences for inferring tumor evolution. Bioinformatics, 34(17):i917–i926, Sep 2018.

[12] Edith M Ross and Florian Markowetz. Onconem: inferring tumor evolution from single-cell sequencing data. Genome Biol, 17:69, Apr 2016.

[13] Hamim Zafar, Anthony Tzen, Nicholas Navin, Ken Chen, and Luay Nakhleh. Sifit: inferring tumor trees from single-cell sequencing data under finite-sites models. Genome Biol, 18(1):178, 09 2017.

[14] Mohammed El-Kebir. SPhyR: tumor phylogeny estimation from single-cell sequencing data under loss and error. Bioinformatics, 34(17):i671–i679, Sep 2018.

[15] R. Schwartz and A. A. Schaffer. The evolution of tumour phylogenetics: principles and practice. Nat. Rev. Genet., 18(4):213–229, 04 2017. [PubMed Central:PMC5886015] [DOI:10.1038/nrg.2016.170] [PubMed:28190876].

[16] Benjamin J Raphael, Jason R Dobson, Layla Oesper, and Fabio Vandin. Identifying driver mutations in sequenced cancer genomes: computational approaches to enable precision medicine. Genome Med, 6(1):5, 2014.

[17] Clemency Jolly and Peter Van Loo. Timing somatic events in the evolution of cancer. Genome Biol, 19(1):95, 07 2018.

[18] Collin M Blakely, Thomas B K Watkins, Wei Wu, Beatrice Gini, Jacob J Chabon, Caroline E McCoach, Nicholas McGranahan, Gareth A Wilson, Nicolai J Birkbak, Victor R Olivas, Julia Rotow, Ashley Maynard, Victoria Wang, Matthew A Gubens, Kimberly C Banks, Richard B Lanman, Aleah F Caulin, John St John, Anibal R Cordero, Petros Giannikopoulos, Andrew D Simmons, Philip C Mack, David R Gandara, Hatim Husain, Robert C Doebele, Jonathan W Riess, Maximilian Diehn, Charles Swanton, and Trever G Bivona. Evolution and clinical impact of co-occurring genetic alterations in advanced-stage EGFR-mutant lung cancers. Nat Genet, 49(12):1693–1704, Dec 2017.

[19] Nabil Amirouchene-Angelozzi, Charles Swanton, and Alberto Bardelli. Tumor evolution as a thera-peutic target. Cancer Discovery, 7(8):805–817, 2017.

[20] Kiya Govek, Camden Sikes, and Layla Oesper. A consensus approach to infer tumor evolutionary histories. In Proceedings of the 2018 ACM International Conference on Bioinformatics, Computational Biology, and Health Informatics, BCB’ 18, pages 63–72, New York, NY, USA, 2018. ACM.

[21] Dikshant Pradhan and Mohammed El-Kebir. On the non-uniqueness of solutions to the perfect phylogeny mixture problem. In RECOMB Int. Conf. on Comparative Genomics, RECOMB-CG’ 18, pages 277–293. Springer, 2018.

[22] Kiran Tomlinson and Layla Oesper. Examining tumor phylogeny inference in noisy sequencing data. In 2018 IEEE Int. Conf. on Bioinformatics and Biomedicine, BIBM’ 18, pages 36–43. IEEE, 2018.

[23] Mohammed El-Kebir, Gryte Satas, Layla Oesper, and Benjamin J Raphael. Inferring the mutational history of a tumor using multi-state perfect phylogeny mixtures. Cell Syst, 3(1):43–53, 07 2016.

[24] Moritz Gerstung, Clemency Jolly, Ignaty Leshchiner, Stefan C Dentro, Santiago Gonzalez Rosado, Daniel Rosebrock, Thomas J Mitchell, Yulia Rubanova, Pavana Anur, Kaixan Yu, et al. The evolutionary history of 2,658 cancers. bioRxiv, page 161562, 2018.

[25] Yusuke Matsui, Atsushi Niida, Ryutaro Uchi, Koshi Mimori, Satoru Miyano, and Teppei Shimamura. phyc: Clustering cancer evolutionary trees. PLoS Comput Biol, 13(5):e1005509, 05 2017.

[26] David F Robinson and Leslie R Foulds. Comparison of phylogenetic trees. Mathematical Biosciences, 53(1-2):131–147, 1981.

[27] Douglas E Critchlow, Dennis K Pearl, and Chunlin Qian. The triples distance for rooted bifurcating phylogenetic trees. Systematic Biology, 45(3):323–334, 1996.

[28] Nikolai Karpov, Salem Malikic, Md. Khaledur Rahman, and S. Cenk Sahinalp. A multi-labeled tree edit distance for comparing “clonal trees” of tumor progression. In Laxmi Parida and Esko Ukkonen, editors, 18th International Workshop on Algorithms in Bioinformatics (WABI 2018), volume 113 of Leibniz International Proceedings in Informatics (LIPIcs), pages 22:1–22:19, Dagstuhl, Germany, 2018. Schloss Dagstuhl-Leibniz-Zentrum fuer Informatik.

[29] Paola Bonizzoni, Simone Ciccolella, Gianluca Della Vedova, and Mauricio Soto Gomez. Does relaxing the infinite sites assumption give better tumor phylogenies? An ILP-based comparative approach. IEEE/ACM Transactions on Computational Biology and Bioinformatics, 2018.

[30] Matt McVicar, Benjamin Sach, Cédric Mesnage, Jefrey Lijffijt, Eirini Spyropoulou, and Tijl De Bie. Sumoted: An intuitive edit distance between rooted unordered uniquely-labelled trees. Pattern Recog-nition Letters, 79:52–59, 2016.

[31] Charles Gawad, Winston Koh, and Stephen R Quake. Dissecting the clonal origins of childhood acute lymphoblastic leukemia by single-cell genomics. Proc Natl Acad Sci USA, 111(50):17947–52, Dec 2014.

[32] El-Kebir Group. OncoLib: Library for tumor heterogeneity, 2018.

[33] Douglas E. Critchlow, Dennis K. Pearl, and Chunlin Qian. The triples distance for rooted bifurcating phylogenetic trees. Systematic Biology, 45(3):323–334, 09 1996.

[34] Peter J. Rousseeuw. Silhouettes: A graphical aid to the interpretation and validation of cluster analysis. Journal of Computational and Applied Mathematics, 20:53–65, 1987.

[35] Salem Malikic, Simone Ciccolella, Farid Rashidi Mehrabadi, Camir Ricketts, Md Khaledur Rahman, Ehsan Haghshenas, Daniel Seidman, Faraz Hach, Iman Hajirasouliha, and S Cenk Sahinalp. Phiscs-a combinatorial approach for sub-perfect tumor phylogeny reconstruction via integrative use of single cell and bulk sequencing data. BioRxiv, page 376996, 2018.

[36] Yong Wang, Jill Waters, Marco L Leung, Anna Unruh, Whijae Roh, Xiuqing Shi, Ken Chen, Paul Scheet, Selina Vattathil, Han Liang, Asha Multani, Hong Zhang, Rui Zhao, Franziska Michor, Funda Meric-Bernstam, and Nicholas E Navin. Clonal evolution in breast cancer revealed by single nucleus genome sequencing. Nature, 512(7513):155–60, Aug 2014.

[37] Peter Eirew, Adi Steif, Jaswinder Khattra, Gavin Ha, Damian Yap, Hossein Farahani, Karen Gelmon, Stephen Chia, Colin Mar, Adrian Wan, Emma Laks, Justina Biele, Karey Shumansky, Jamie Rosner, Andrew McPherson, Cydney Nielsen, Andrew J. L. Roth, Calvin Lefebvre, Ali Bashashati, Camila de Souza, Celia Siu, Radhouane Aniba, Jazmine Brimhall, Arusha Oloumi, Tomo Osako, Alejandra Bruna, Jose L. Sandoval, Teresa Algara, Wendy Greenwood, Kaston Leung, Hongwei Cheng, Hui Xue, Yuzhuo Wang, Dong Lin, Andrew J. Mungall, Richard Moore, Yongjun Zhao, Julie Lorette, Long Nguyen, David Huntsman, Connie J. Eaves, Carl Hansen, Marco A. Marra, Carlos Caldas, Sohrab P. Shah, and Samuel Aparicio. Dynamics of genomic clones in breast cancer patient xenografts at single-cell resolution. Nature, 518:422, November 2014.

